# Application of spatial transcriptomics across organoids: a high-resolution spatial whole-transcriptome benchmarking dataset

**DOI:** 10.1101/2025.05.04.651803

**Authors:** MR Nucera, N Charitakis, RF Leung, A Leichter, N Tuano, M Walkiewicz, V Sawant, L Rowley, M Scurr, PX Er, KS Tan, R Sutton, F Ahmad, R Saxena, A Maytum, DL Turner, HK Voges, HT Nim, X Sun, B Yang, KL Li, G Ball, AG Elefanty, SR Lamandé, KT Lawlor, JM Vanslambrouck, RJ Mills, ES Ng, EG Stanley, RB Werder, MH Little, DA Elliott, ER Porrello, MC Faux, DD Eisenstat, S Velasco, FJ Rossello, M Ramialison

## Abstract

Stem cell-derived organoids hold promise to model tissue-specific disease. To enable this, it is crucial to assess how transcriptional signatures, cellular organisation and composition of organoids compare to *in vivo* counterparts. However, technologies which elucidate regional molecular identity, like spatial transcriptomics, have been challenging to apply to organoids. This study presents the first systematic profiling of multiple stem cell derived organoid models (brain, heart muscle, heart valve, kidney, lung, cartilage, and haematopoietic) with Stereo-seq, a full transcriptome, spatial transcriptomics assay using on-chip in situ RNA capture at subcellular resolution. It describes optimisation of this assay to characterise organoids, use of multiple organoid samples on a single chip, assess differences in RNA capture efficiency compared to reference tissues and its limitations. This study introduces a bespoke analysis method that partitions samples into regions and further characterises them. These findings inform future works to characterise organoids using spatial transcriptomics, providing insights in optimising RNA capture of multiple organoids across a chip and novel methods for regional analysis.

## Introduction

Pluripotent stem cell (PSC)-derived organoid models are used to investigate human development and disease pathogenesis and provide a platform to measure drug response [1]. This requires organoids to accurately recapitulate the corresponding in vivo tissue, both physiologically and in cellular composition, by replicating in vitro the process of native stem cell development and organization [2], [3].

Spatial transcriptomics (ST) technologies capture gene expression across tissues while maintaining anatomical and morphological structure. ST encompasses a wide array of technologies that are divided into imaging- and sequencing-based assays [4]. This study focuses on the application of Stereo-seq, a sequencing-based technology capable of capturing transcriptome-wide gene expression at single cell resolution [5]. Stereo-seq profiles an embedded tissue section using a specialised chip, that simultaneously allows mRNA capture and imaging. Given the high cost of ST experiments per sample, finding a cost-effective approach to profile multiple organoids using a single chip is critical. While ST has previously been applied to a single type of organoid being placed on Stereo-seq chip [6], [7], [8], technical challenges arise when using small size organoids that cover a limited portion of the ST chip. This may result in artificially lower mRNA levels or capture of fewer gene transcripts, defined throughout this manuscript as the total data capture. To date, there are no studies that directly compare mRNA expression and gene profiling of organoids against in vivo tissue. Furthermore, the placement of several individual organoids on a single chip has proven to be viable for certain organoid types [8], yet studies describing the feasibility and benefits of simultaneous placement of multiple organoid conditions on the same chip is limited.

This study performs a systematic spatial transcriptome profiling of multiple organoid models (brain, heart muscle, heart valve, kidney, lung, cartilage, and haematopoietic) using the Stereo-seq platform compared to two in vivo reference E13.5 mouse head and 21 post-conception week (PCW) mouse heart tissues. The study describes the successful placement of multiple organoids of the same tissue type - cultured under different conditions for certain tissues - on the same chip. Subsequently, the platform’s capability to clearly distinguish transcriptional gene signatures from these different individual organoids on the same Stereo-seq chip was tested. The effect of size, shape, and cell-type composition of different organoids on data capture, when compared to one another and to larger, in vivo reference E13.5 mouse head and 21 PCW mouse heart tissues was investigated. Finally, despite challenges in cell type annotation linked to challenges in achieving single-cell resolution, this study illustrates an effective analysis for uncovering region-specific differences across organoids as an alternative to cell clustering and classification. This approach enables the identification of spatially resolved biologically relevant gene expression patterns across multiple organoid models for the first time.

## Results

This study systematically profiled the transcriptome of eight human organoid models (brain, lung, haematopoietic, transwell kidney, suspension kidney, heart valve, cartilage, and heart muscle) using the Stereo-seq platform. A modified Stereo-seq workflow was designed to capture multiple conditions where multiple organoids of the same tissue type were co-embedded and placed on a single Stereo-seq chip (see Methods) (Fig. 1A). To prevent tissue detachment from the chip surface, observed in cartilage, heart muscle, and kidney organoids, Stereo-seq chips were coated with poly-L-lysine, as part of our modified Stereo-seq workflow. To limit transcript diffusion and allow for accurate spatial gene expression profiling, tissue-specific cell permeabilization also required optimisation. This optimization was critical to allow processing of multiple organoid samples, derived from the same tissue, and different culture conditions on the same chip. Despite that suitable permeabilization conditions could be identified for most models, haematopoietic and suspension kidney organoid samples displayed higher levels of transcript diffusion, revealed by transcripts captured outside of tissue locations, despite optimised permeabilization conditions (Fig. 1A).

**Figure 1 –.**
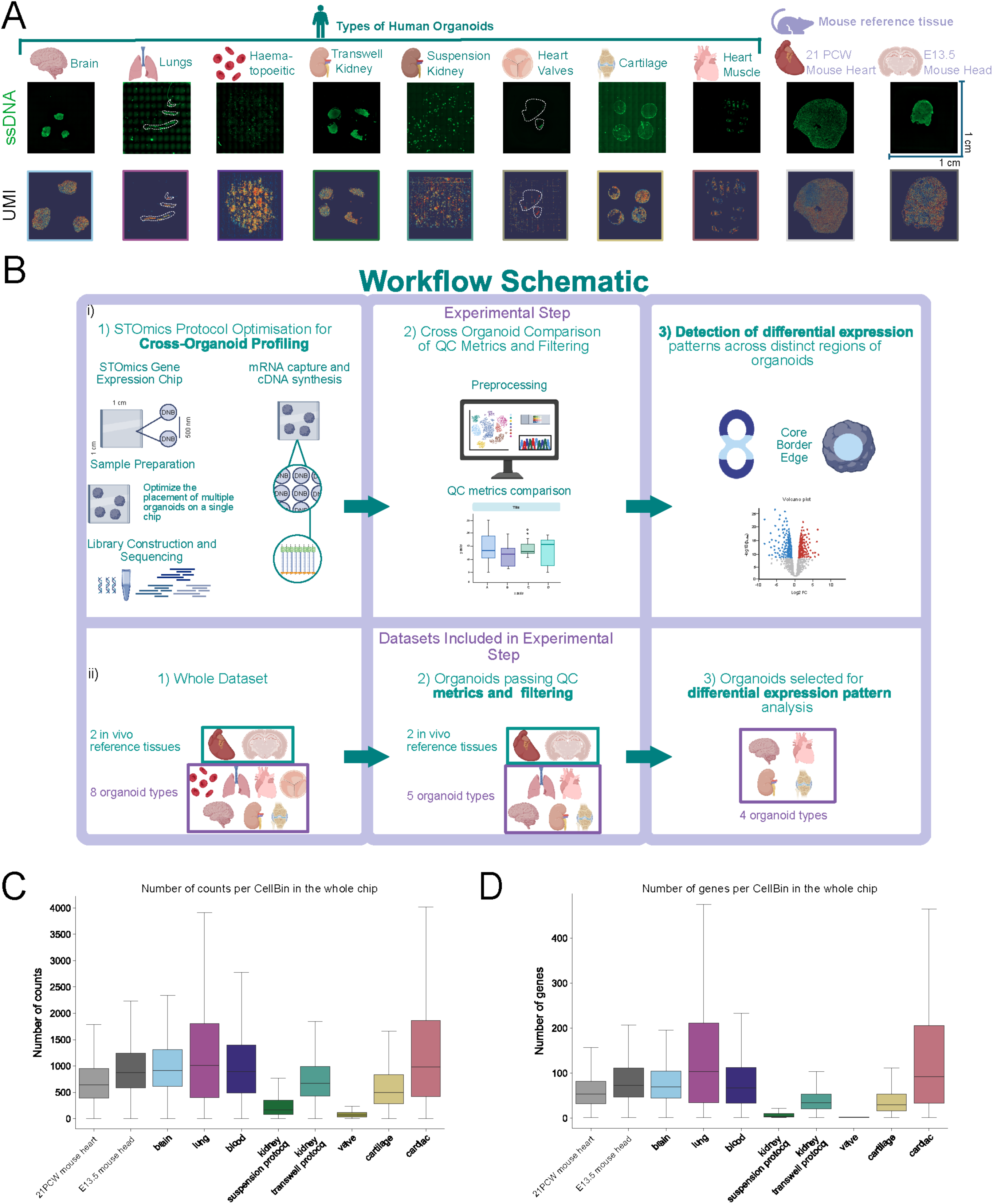
Overview of quality metrics across multiple organoid types and in vivo reference tissues when captured using the Stereo-seq platform. (A) Schematic of organoid models and in vivo reference tissues included on individual Stereo-seq 1cm2 chips. Immunofluorescence images display regions of ssDNA capture across the chip while accompanying areas of high-count expression are visualised as UMI heatmaps. White dotted lines indicate organoid boundaries on UMI heatmaps and immunofluorescence images. (B) Schematic of experimental design. i) Steps included from sample preparation and optimisation through to data analysis and investigation of regional patterns. ii) Outline of datasets included in each experimental step. Protocol optimisation and cross-organoid profiling was performed on a total of 10 Stereo-seq chips, divided across two in vivo reference tissues and eight organoid types. After assessing chips that passed quality control filters, both in vivo reference tissues and five organoid types were retained for analysis. Finally, four organoid models were selected for region-specific analysis patterns of gene expression. (C) Number of UMI counts per CellBin across each chip. (D) Number of genes per Cellbin across each chip.

Two reference tissues (E13.5 mouse head and 21 PCW mouse heart) were also profiled using the standard Stereo-seq workflow, with the aim to use these datasets as baselines against the characterised organoid models (Fig. 1A). To account for the custom Stereo-seq workflow and compensate for differences in RNA capture across organoids, a custom bioinformatics workflow was developed, including a region-specific analysis, described in detail below (Fig. 1B). This workflow filtered out low quality organoid samples to ensure comparisons to baseline were only drawn for models for which the protocol optimisation had been successful. This then allowed investigation of the increased chip coverage on data capture, to determine compatibility of the Stereo-seq platform with multiple organoid models. RNA capture quality across the entire chip was then assessed through a variety of metrics: comparison of the number of captured counts and genes on a per CellBin basis (Fig. S1C-S1D), captured DNA nanoballs and unique reads (Fig. S1A-B) and sequencing saturation (Fig. S1C). Firstly, data capture was evaluated on CellBins (equivalent to single cells visualised across the corresponding immunofluorescence ssDNA image), and the smallest analytical unit generated during pre-processing via default cell-segmentation methods on the accompanying immunofluorescence image (ssDNA, Fig. 1A). Although large variation persisted in total number of transcripts captured between organoids and in vivo tissue reference (Fig. S1A-S1B), capture range for both number of genes and counts per CellBin were found to be comparable across organoid models and in vivo reference tissues (Fig. 1C-1D), irrespective of sequencing saturation. It is worth noting that only kidney suspension organoids did not reach a saturation of 80% (Fig. S1C). This suggests that the observed variation could have been introduced by differences in chip coverage area, rather than organoid type, emphasised by a positive correlation between total UMIs passing QC and tissue size of organoid samples compared to in vivo reference tissue (Fig. S1D).

The heart valve and kidney suspension protocol models showed a reduced RNA capture relative to other organoid models, evidenced by a low number of DNA nanoballs captured under tissue along with low numbers of mapped unique reads (Fig. S1A-S1B). Additionally, the haematopoietic organoid chips, displayed an unexpectedly broad data capture range (Fig. S1A-S1B). This was likely due to the placement of a large number of organoids per chip leading to increased transcript diffusion, mainly evidenced by the presence of UMIs outside tissue regions of ssDNA (Fig. 1A). Given these issues, downstream analyses focused on organoid models with uniform size and shape, resulting in an increased RNA capture across chips. Individual brain, transwell kidney, and cartilage organoid samples placed on the same chip showed comparable detected number of genes and transcripts captured compared to E13.5 mouse head and 21 PCW mouse heart in vivo references (Fig. S1E-S1I). For quality assurance, the percentage of CellBins expressing housekeeping genes GAPDH/Gapdh and ACTB/Actb across in vivo references and individual brain and cartilage organoid samples were investigated (Fig. S2A). While cell-to-cell expression levels of housekeeping genes varied (Fig. S2C), similar to previously reported single-cell RNA-Seq experiments [9], no clear spatial expression patterns of GAPDH/Gapdh and ACTB/Actb across the organoids passing QC filtering (Fig. S2A-S2C) were observed, reflecting the expected homogenous expression nature of these genes. These observations also revealed no systematic differences introduced by coating chips with poly-L-lysine (Fig. S2C).

In addition to CellBin segmentation, Stereo-seq experiments can also be analysed using a region-based approach (“square bin”) by grouping adjacent transcript capture areas together. Given the smaller size of the organoids compared to mouse reference tissues and RNA capture variability across all profiled tissues (Fig. 1), the effect of increasing bin size on capture of RNA levels, genes, and biological signal was explored next (Fig. 2). The effects of altering square bin size, which determines the resolution of defined regions, revealed loss of tissue coverage at larger bin sizes, which could further reveal differences in platform performance when profiling in vivo reference tissues and organoid models (Fig. 2A-D). For both in vivo references and different organoid models, when the bin size is increased from 50 to 75, the expected accompanying proportional decrease in the number of bins passing QC was not observed (Fig. 2Ei-ii).

**Figure 2 –.**
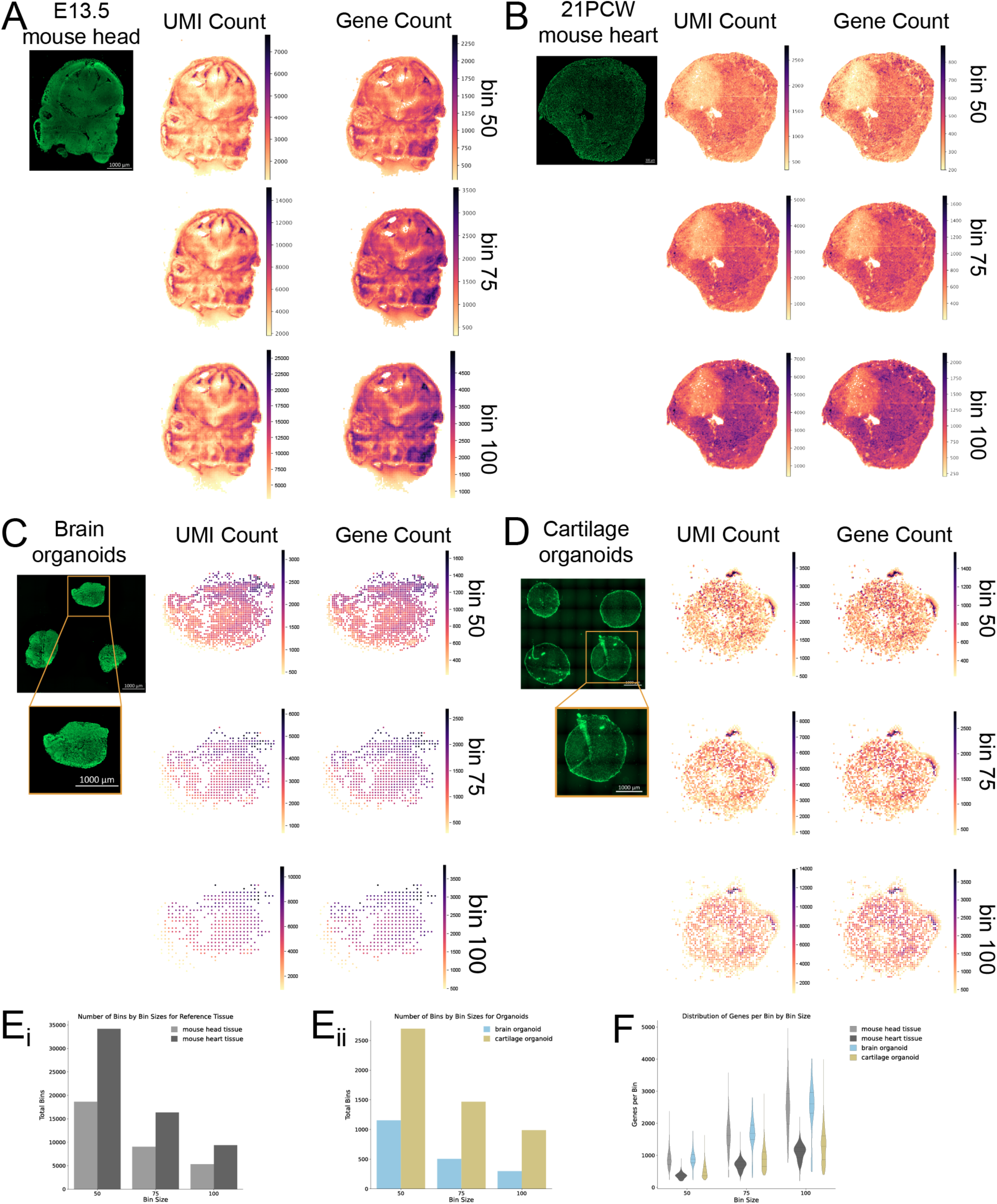
Effect of increasing bin size (50, 75, 100) on spatial patterning and data visualisation across in vivo reference tissues and organoids. (A) Immunofluorescence image of ssDNA capture and heatmaps visualising regions of high counts and genes per bin across in vivo reference tissue E13.5 mouse head at increasing bin sizes. (B) Immunofluorescence image of ssDNA capture and heatmaps visualising regions of counts and genes per bin across in vivo reference tissue 21 PCW mouse heart tissue at increasing bin sizes. (C) Immunofluorescence image of ssDNA capture and heatmaps visualising regions of counts and genes per bin across a single brain organoid at increasing bin sizes. Brain organoid 1 displayed in the heatmaps is highlighted in immunofluorescence image. (D) Immunofluorescence image of ssDNA capture and heatmaps visualising regions of counts and genes per bin across a single pseudo-replicate of cartilage organoid. Cartilage organoid 1 displayed in the heatmaps is highlighted in immunofluorescence image. (E) Bar plots of total number of bins captured across tissue at different bin sizes. (i) In vivo reference E13.5 mouse head and 21 PCW mouse heart tissue. (ii) Brain organoid 1 and Cartilage organoid 1. (F) Distribution of number of genes captured in each bin at bin sizes 50, 75 and, 100 for in vivo reference E13.5 mouse head, 21 PCW mouse heart, brain organoid 1 and cartilage organoid 1.

Our analysis demonstrated that increasing bin size from 50 to 75 to 100 (25 μm^2^, 37.5 μm^2^ and 50 μm^2^ respectively) accompanied by analysis using bespoke filtering parameters to eliminate inclusion of background signal changes the overall tissue coverage area in both the in vivo reference tissues and brain and cartilage organoids at larger bin sizes (Fig. 2, Table S1). However, due to the reduced size of organoids samples in general, this loss of largely peripheral signal may contribute to downstream challenges to identify cell type compositions in the organoids.

It is expected that a bin size increase of 50% would produce a 50% decrease in the number of bins. In both in vivo reference tissues, increasing the bin size by 50% led to the number of bins decreasing by more than half, while a doubling of bin size leads to only about one-third of the total number of bins being retained (Fig. 2Ei, Table S1). In mouse head tissue this created a drop from 18634 bins at bin size 50 to 9024 bins at bin size 75 and 5313 at bin size 100 (Table S1). This disproportionate reduction in the number of bins with increasing bin size was more noticeable across organoids compared to in vivo reference tissues (Fig. 2Eii, Table S1). For the brain organoids, bin size 50 results in a tissue area of 67450 μm^2^ being analysed, and this fell to 49350 μm^2^ with analysis performed at bin size 100 (Table S1). We also found that larger bin sizes necessitated stricter filtering parameters to eliminate the inclusion of background signal in the tissue region, leading to removal of certain tissue areas for downstream analysis and resulting in fewer bins (Table S1). These asymmetrical changes were mirrored in the total calculated tissue area at different bin sizes for both in vivo reference tissues and organoid models (Table S1). In addition, increasing the bin size across organoid and reference tissues led to changes in the range of number of genes captured per bin (Fig. 2F), with the range of distribution number of captured genes increasing as bin size increased.

Overall, RNA capture proved more effective at smaller bin sizes while minimising background signal across the aforementioned tissues. Additionally, smaller bin sizes allowed for maximum tissue area to be included in downstream analysis without the addition of background noise to the dataset in both in vivo reference and organoid tissues. However, the relatively low number of captured UMIs and genes across individual samples, either with CellBins or a square bin size of 50 above (Fig. 1C-1D), hindered analysis at a single-cell resolution using this platform (Fig. 2C-2D).

To further investigate the feasibility of multi-sample placement on single chips, the effect of normalising samples across the entire chip was compared to a sample-specific normalisation, performed independently on samples after sub-setting (Fig. S3). Overall, the spatial expression of housekeeping (GAPDH, ACTB) or organoid marker genes remained unaffected across both normalisation methods (Fig. S3). Evidence that normalisation did not skew biological signals is reflected in the expected homogeneous expression of COL2A1, COL9A3 and SPARC across cartilage organoids, given that they are composed almost exclusively of chondrocyte cells (Fig. S3).

Despite increasing bin sizes and normalisation, data generated from organoid experiments did not allow for cell clustering and further cell type classification primarily due to insufficient capture of cell type defining transcripts and genes. This was contrasted with the clear and accurate spatial profiles of the in vivo samples (Fig. S2B-S2D). Therefore, an alternative, exploratory analysis method that considers organoid regions was developed. This approach allows comparison of gene expression profiles between border and core regions of organoids, a valuable analysis given previous findings observing region-specific transcriptional differences within organoids [1], [10], [11] (Fig. 3A)

**Figure 3 –.**
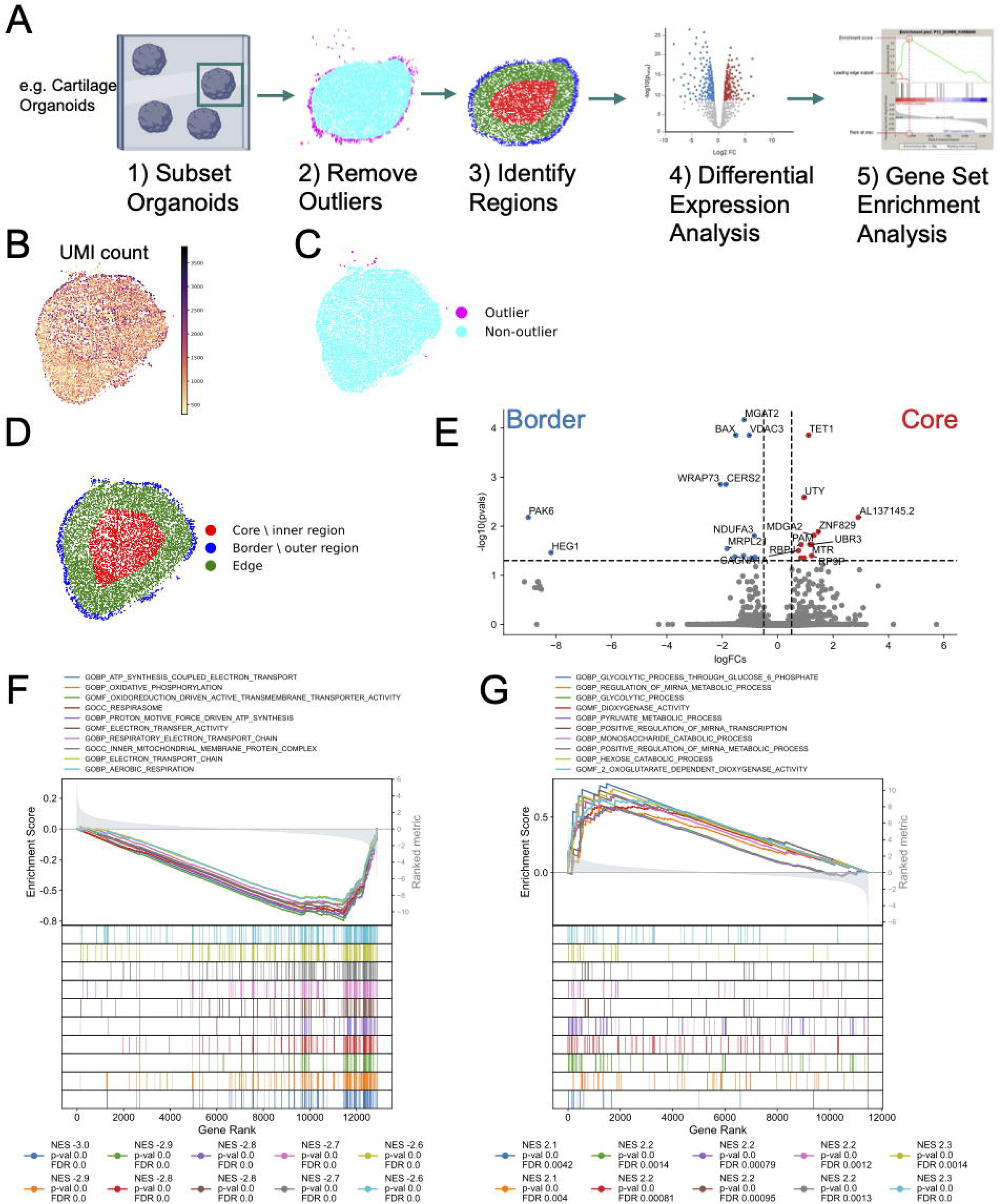
Analysis of spatially-restricted, differentially expressed genes across centre and border regions. (A) Schematic of representative workflow of region-specific analysis between core and border regions demonstrated using cartilage organoids. (B) Heatmap of regions of UMI counts across brain organoid 1 using CellBin. (C) Outlier CellBins identified and visualised in magenta across brain organoid 1, representing a false positive signal to be removed before region analysis. (D) Edge (green), border (blue) and centre (red) regions highlighted for analysis in brain organoid 1. (E) Volcano plots of genes upregulated in core (visualised in red) and those upregulated in border region (visualised in blue) for brain organoid pseudo-replicates 1-4. (F) Gene set enrichment analysis output showing top 10 negatively enriched terms in core of brain organoids. Each curve represents the enrichment profile the gene set. Legend annotations include Normalised Enrichment Score, p-value, and FDR q-value. (G) Gene set enrichment analysis output showing top 10 positively enriched terms in core of brain organoids.

After removing outliers (bins falling outside of the organoid boundaries), our approach classifies remaining bins into inner and outer regions, based on their distance from the centre of the organoid (see methods). Gene expression from bins within each region is then aggregated using a pseudo-bulk strategy, resulting in a stronger signal that overcomes the low expression of individual bins. We selected brain, transwell kidney, and cartilage organoids for this analysis, based on comparable shape and size of organoid samples of the same tissue type, which allowed benchmarking of the developed region-specific analysis method (Fig. 3 and Fig. S4A-S4I).

The initial quality-filtering step measured the distance from each bin located at the extremes of the organoid boundaries to its k nearest neighbouring bins and removed those that were isolated from the rest. This method successfully excluded any low-quality regions (based on UMI counts) that potentially arose from inconsistent tissue adherence to the chip (Fig. 3A-C, Fig. S4A-S4F). From this analysis, brain and cartilage organoid samples exhibited fewer outliers than kidney, likely due to their uniform shape and lesser number of detached cells at the edge of the organoids (Fig S4D-S4F). This analysis subsequently enabled the identification of edge, border, and core regions within the assessed organoid models (Fig. 3D, Fig. S4G-S4I). A differential expression analysis between core and border regions of organoid models, using a pseudo-bulk analysis and organoid samples as replicates, followed. It should be noted that this approach does not account for spatially variable gene expression. In cartilage organoids, no differentially expressed genes (DEGs) were identified between these regions, confirming the homogenous cell population expected in this model [12] (Fig. S4J). In contrast, brain organoids showed transcriptional differences between border or core regions, with 13 genes differentially upregulated in core and 13 genes in border regions (Fig. 3E). Gene set enrichment analysis (GSEA) of biological processes showed that border regions of these organoids were enriched for gene sets related to ATP synthesis and electron transport (Fig. 3F), whereas core regions were enriched for gene sets related to glycolysis (Fig. 3G). These observations align with previous studies showing enrichment of glycolytic gene sets in cortical neural progenitors which are located at the core of brain organoid [13], while more energy-demanding neuronal populations are found at the borders of organoids at this developmental stage [13]. These findings demonstrate that the proposed region-specific approach allowed the identification of spatially and metabolically distinct cell populations in brain organoid models, which was also observed when using a Slide-seq assay, an alternative sequencing ST technique [13].

We further expanded the use of our method to on irregularly shaped heart muscle organoids. These organoids have a “figure eight” shape, with two pole regions on either side of the organoid and a centre region, amenable to measurement contractile functions (Fig. 4A-4B) [14]. Heart muscle organoids were cultured under two different conditions, using a previously published standard serum-free method (control) [15] and a directed maturation method (DM) for enhanced maturation and contractile force [16] (Fig. 4A-4B). By applying the developed region-specific analysis, pole regions were filtered out from downstream analysis due to low cell density and mRNA capture (Fig. 4A-4B). This shows that this method could act as a secondary quality control based on tissue distribution. Centre regions of the control and DM organoids were compared using organoid samples as replicates (Fig. 4B, Fig. S4K-S4L), showing transcriptional differences between the compared conditions (Fig. 4C). This analysis identified an upregulation of MRPS12, a mitochondrial riboprotein, in the DM-cultured organoids (Fig. 4D), suggesting a difference in mitochondrial activity in organoids culture in DM-media. In control heart muscle organoids, a GSEA analysis identified enriched gene sets implicated in ion channel activities (Fig. 4E), while gene sets linked to mitochondrial activities, ATP synthesis, and metabolism were upregulated in DM heart muscle organoids (Fig. 4F). Enrichment of these gene sets reflect recent findings that reported alternative usage of ion channels and increase in sarcomere maturity of organoids cultured in DM media indicated through the downregulation of ion channel genes and upregulation of metabolic genes, respectively [16]. The transcriptional differences captured through ST between serum-free control and DM heart muscle organoids resemble previous characterisation of these models, demonstrating the validity of ST assays to capture transcriptional differences of organoids grown under different conditions.

**Figure 4 –.**
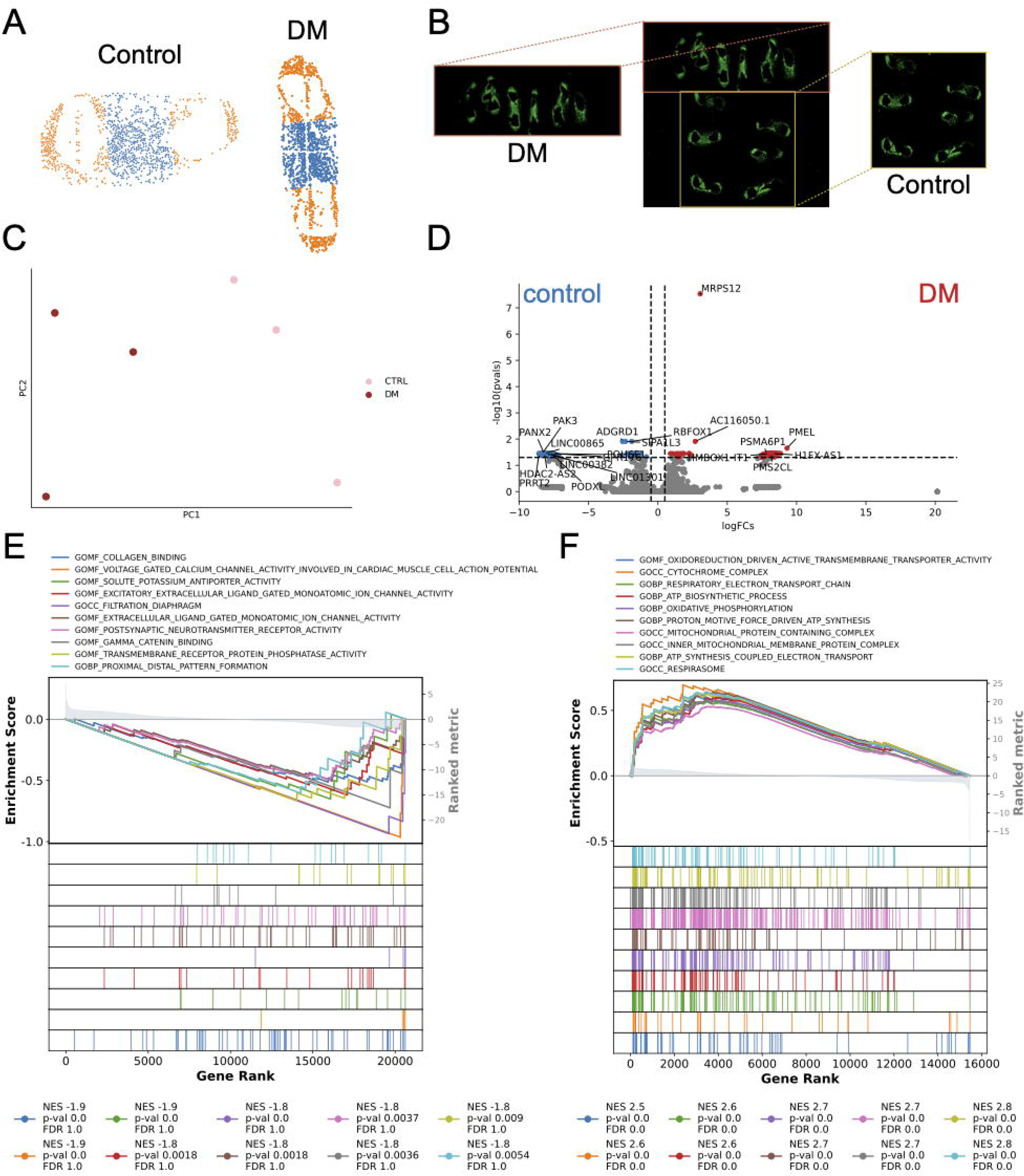
Analysis of differentially expressed genes across centre regions of heart muscle organoids of different conditions. (A) Example of outlined centre (blue) and pole (orange) regions across DM and control heart muscle organoids. (B) Overview of immunofluorescent heart muscle organoids across Stereo-seq chip outlining placement of six DM organoids on the top and six control organoids on the bottom side of the chip. (C) PCA plot of data from 3 centre regions of control organoids and 3 regions of DM organoids. (D) Volcano plots of genes upregulated in centre of 3 DM organoids (red) compared to genes upregulated in centre of 3 control organoids (blue). (E) Gene set enrichment analysis showing top enriched terms in the centre of control heart muscle organoids. (G) Gene set enrichment analysis showing top enriched terms in the centre of DM heart muscle organoids.

## Discussion

This study assessed the performance of Stereo-seq, a ST by sequencing assay, in eight iPSC-derived human organoid models and two mouse in vivo tissue references. The main objective was to validate this technology across multiple organoid tissue types in comparison to reference tissues, in which spatial gene expression profiles showed anatomical specificity at single cell resolution.

Across the eight organoid models assessed, cartilage, brain, and transwell kidney organoids resulted in sample quality comparable to tissue references, which could be attributed to their size and relatively uniform, round shapes. Despite this, RNA capture of these samples failed to achieve single cell gene expression resolution. Organoids with smaller surface areas, such as suspension kidney and haematopoietic organoids (notwithstanding comparable transcript capture and total gene counts) displayed high amounts of diffuse gene expression captured outside of the organoid surfaces. We suspect this was likely due to additional challenges presented by these organoids, cultured in more of a ‘liquid-phase’ compared to other tissue types, resulting in transcript diffusion. Another challenge in applying Stereo-seq to these organoids was their lack of adhesiveness, likely contributing to non-uniform transcript capture across chips. As a remedy, poly-L-lysine chip coating was used to maximise tissue adhesion (see Methods). The data generated from heart muscle organoids demonstrated that this method was effective in improving organoid adhesion and should be considered for all organoids to improve data quality.

This study tested the feasibility of placing multi-sample placement on a single chip. Previously, ST technology required single tissue sections to be placed on separate chips to yield optimal outcomes [6], [7], [8]. Our analysis revealed that placing up to 12 sections on the same chip, as in the case of heart muscle organoids, still resulted in accurate transcript capture, both cultured in the same media, and organoids grown in different culture media, such as the case of control vs DM heart muscle organoids. The ability to maximise coverage with multiple sections per chip greatly extends the potential use of ST technology for organoids, ensuring high data value per chip and reduce inter-chip variability.

The expression of housekeeping genes such as GAPDH and ACTB was uniform across cartilage, brain, and transwell kidney organoids, although only in less than 50% of the identified CellBins, resembling expression patterns of these genes in in vivo reference tissues. However, the identification of region-specific transcriptional signatures in organoids was limited when using CellBins (i.e. single-cell resolution), likely due to the relatively low number of transcripts and genes captured per CellBin. While the number of transcripts and genes detected per bin were comparable to other studies utilising Stereo-seq [5], it was found to be lower than the accepted range for single-cell RNA-seq experiments. Interestingly, increasing square bin sizes, at the expense of losing single cell resolution, did not resolve the challenges in detecting region-specific expression patterns in organoids models. This may also be a product of the unbiased transcript capture technology coupled with low number of transcripts captured, leading to an overrepresentation of abundant transcripts. This shows that while whole-transcriptome ST, including Stereo-seq and Visium HD, could capture transcripts from organoid models, its utility for identifying region-specific expression patterns at single cell level in these model systems was limited. However, this higher-throughput approach for characterisation of organoids could be useful for discerning spatially distinct molecular signatures, as previously shown to be viable using in vivo tissue samples signatures [5], [17].

Given the smaller size of organoids compared to in vivo tissue, the structural compartments in these models could be relatively subtle and marked by fewer cells. Coupled with the challenge of achieving single-cell resolution, this hindered the ability of ST technique in capturing spatially resolved gene expression in organoids and hence identification of defined patterning. The use of cryo-sections may also limit the assessment of these organoids, as structural compartments could be rare and difficult to capture on a single cell-thick layer. These challenges demonstrate the need to further optimise protocols to apply ST to organoid models. As mentioned above, improvements could include applying alternative chip coating methods to further increase tissue adhesion or organoid fixation to better preserve spatial gene expression profiles.

As standard clustering and cell type classification approaches could not be applied to dataset from organoids of this study, we developed an alternative, exploratory regional analysis method that allowed spatial gene expression profiling in organoids between core and border regions. We demonstrated its power in identifying region-specific biological processes as well as its utility as a secondary quality control filter when applied to brain and heart muscle organoids. Given the use of Euclidean distance to calculate outliers, the application of this analysis as a secondary quality control filter to round organoids enables more accurate calculation of cells that have detached from main organoid bodies. As mentioned, the ability to understand region-specific gene expression patterns in these models could be key to elucidating general structural organisation that may be present in these systems. This knowledge could then be used to further advance organoid models to better reflect their in vivo counterparts, especially with regards to progressing organoid patterning for the study of developmental tissue patterning. This could have significant implications in developmental biology, such as identifying processes of cell differentiation. As ST platforms for RNA capture continue to evolve and expand imaging options, an updated Stereo-seq protocol incorporating H&E imaging, a morphology-based analysis could be incorporated into future analytical pipelines.

Overall, this study explores the use of profiling multiple organoid types using ST platform. It demonstrates the feasibility of placing multiple organoids generated under different culture conditions on a single Stereo-seq chip, allowing capture of distinct biological signatures from each model. Furthermore, we illustrate the abilities and limitations of profiling a range of organoid types with the current Stereo-seq platform, identifying potential steps for future optimisation such as use of poly-L-lysine for adhesion. Given the limitation around current RNA capture in allowing organoid-specific cell-type annotation, which is not mitigated by increasing bin size for analysis, we developed a regional-specific analysis workflow for capturing biologically relevant gene expression. This provides a basis for future optimisation of organoid profiling using the Stereo-seq platform, including maximising data capture from a single experiment.

## Limitations of Study

Although the application of Stereo-seq ST technology across a wide range of organoids was examined, other ST technologies may present different outcomes when applied to organoid systems. For instance, other sequencing-based ST technology such as 10x Visium HD, also offers spatial gene expression profiling at single cell resolution, and, given the different chemistry and transcript capture technology, it could produce alternative results [18].

Alternatively, probe-based imaging ST technologies could also perform differently, with sub-cellular resolution and higher specificity. To validate findings reported in this study, traditionally spatial gene expression profiling methods such in situ hybridisation or immunofluorescence could be performed to assess the accuracy of the gene expression patterns identified. Additionally, expanding the range of in vivo tissues to correspond to the organoid systems included, such as including mouse kidney and lung tissues, would provide additional points of reference to validate the use of Stereo-seq ST technology on various organ and organoid systems.

## Methods

### Animal maintenance and ethics approval

All animals were housed in the Murdoch Children’s Research Institute (MCRI) Disease Model Unit and all animal works performed were conducted under the approval and in compliance of the MCRI Animal Ethics Committee agreements (SPPL20327). Timed pregnant mice were sacrificed by cervical dislocation following MCRI AEC approved protocols at embryonic day 13.5 where the presence of a vaginal plug after mating indicated (E0.5). E13.5 mouse heads and 21 postconceptional week (PCW) mouse hearts were fresh frozen in OCT (SAKURA Cat# 4583) and stored at −800C until further processing.

### Maintenance of induced pluripotent stem cells (iPSC)

PB001.1 iPSCs (male) [19] were used for haematopoietic organoid differentiation, CRL1502.2 iPSCs (female) [20] were used for transwell kidney organoid differentiation, MAFBmTagBFP:GATA3mCherry reporter iPSCs [20] were used for suspension kidney organoid differentiation, PB10.5 iPSCs (male) [19] were used for heart organoid differentiation. Brain organoids were generated from PB001.1 [19] PB001.1 harbouring a heterozygous 1bp insertion at chr5:177,135,105 (c.2insG) and PB001.1 harbouring a homozygous 4bp deletion at chr5:177,135,106 (c.3_6delGGAT). BU3 NGST CRISPRi iPSCs (male) [21] were used for lung organoid differentiation. MCRIi019-A iPSCs (female) [22] were used for cartilage organoid differentiation. Mycoplasma contamination in all iPSCs were monitored and excluded by regular testing. Karyotyping of all iPSCs were performed regularly and no significant genomic imbalances were identified.

All iPSCs, with the exception of those used for brain organoids were grown on human embryonic stem cell (hESC)-qualified Matrigel Matrix (Corning Cat# 354277), with varying medium conditions. iPSCs for brain organoids were grown on Geltrex™ LDEV-Free Reduced Growth Factor Basement Membrane Matrix (Thermo Fisher Scientific Cat # A1413301). iPSCs used for differentiation of cartilage, transwell kidney, suspension kidney, and haematopoietic organoids were maintained in Essential 8 Medium (Thermo fisher Scientific Cat# A1517001). iPSCs used for lung organoid differentiations were maintained in StemFlex Medium (Thermo Fisher Scientific Cat# A3349401), whereas iPSCs for brain and heart organoid differentiation were maintained in mTESR1 Medium (Stem Cell Technologies Cat# 85857).

### Organoid preparation for Stereo-seq

Eight organoid models were generated in accordance to previously described differentiation protocols, including lung airway and alveolar type 2 organoids [23], [24], [25] haematopoietic organoids [26], heart organoids serum-free controls [14], [16], [27], heart organoids with directed maturation (DM) [28], cartilage organoids [12], heart valve organoids [27], transwell kidney organoids [28], suspension kidney organoids [29], and brain organoids [13], [30]. Specifically, following alterations were included in respective organoids to generate different experimental conditions. Both iPSC-derived lung organoids at air-liquid interface were infected on the apical surface with respiratory syncytial virus (RSV-A2, MOI 1) as previously described for SARS-CoV-2 [31]. After 4 hours, the inoculum was removed and cells washed. Uninfected and infected samples were collected 72 hours post infection. From day 48, cartilage organoids were either grown without additional treatment for 14 days or treated as follows. Condition 1: cartilage organoids were cultured without additional treatment after day 48 for 7 days followed by addition of 10nM Triiodothyronine (Sigma Aldrich Cat# T6397) for 7 days, condition 2: cartilage organoids were cultured without additional treatment after day 48 for 7 days, followed by 7 days in osteogenic medium [12] and condition 3: cartilage organoids were grown with addition of 10nM Triiodothyronine for 7 days followed by 7 days in osteogenic medium. All cartilage organoid conditions were harvested at day 62. Brain organoids grown from all three cell lines were collected on day 31.

At the time of harvest, organoids were separated from culturing media and washed in PBS, then placed in pre-chilled OCT. Organoids were then transferred to plastic mould containing OCT, and frozen in an ethanol or isopropanol slurry. For brain organoids, three organoids were embedded together and placed on the chip, whereas four transwell kidney, and cartilage organoids were embedded together and placed on the same chip. Eighteen heart valve organoids were embedded together and placed on the same chip. Six heart organoids of each condition (serum-free and DM) were embedded together in an array of 12 organoids. Over 20 haematopoietic and suspension kidney organoids were embedded in a single mould. For lung organoids, a pair of airway and alveolar type 2 sections, either infected or non-infected, were embedded in OCT together. However, the uninfected airway section was lost during sample processing, resulting in only infected airway section present and alveolar type 2 sections remained.

### Stereo-seq T-chip coating

For the heart muscle, heart valve, and cartilage organoids, the sections were mounted on poly-L-lysine coated Stereo-seq T-chip to enhance adhesion. Stereo-seq T-chips were washed using nuclease-free H2O twice and dried, then coated with 100µL of 0.01% (w/v) poly-L-lysine (Sigma Aldrich Cat# P8920) for 10 minutes at room temperature. Post incubation, the chips were washed with nuclease-free H2O twice, and dried on a 37oC slide dryer.

### Stereo-seq sample processing and library preparation

Samples were prepared and processed using the STOmics Transcriptomics kit (STOmics Cat# 111KT114) in accordance with manufacturer’s instructions (BGI Stereo-seq Transcriptomics set for Chip-on-a-slide user manual version B) with slight variations. 10LJm sections of mouse tissue samples including E13.5 heads and 21PCW hearts, brain organoids, kidney organoids, lung organoids, and haematopoietic organoids were cryo-sectioned on CM3050S cryostat (Leica) and mounted on room temperature T-chips, then processed according to the STOmics user manual. 10LJm cryo-sections of cartilage, heart muscle, and heart valve organoids were placed on room temperature poly-L-lysine coated Stereo-seq T-chips, then processed according to the STOmics user manual. All cryo-sections were fixed in ice cold methanol for 30 minutes at −30oC. Tissue section permeabilization times were as follows: lung, haematopoietic, transwell kidney, suspension kidney, and cartilage organoid sections were permeabilised for 12 minutes, mouse head and brain organoids for 15 minutes, heart organoids and engineered heart valves for 18 minutes, and mouse heart for 24 minutes. The cDNA purification and library preparation of each sample was constructed using the STOmics library preparation kit (STOmics Cat# 111KL114) in accordance with manufacturer’s instructions (BGI STOmics Stereo-seq Transcriptomics set for Chip-on-a-slide user manual version B). Quality of the final cDNA library size was assessed using D1000 Tapestation Reagent (Agilent Technologies Cat# 5067-5602) and Screentape (Agilent Technologies Cat# 5067-5582) and the concentration measured with Qubit dsDNA HS kit (Thermo Fisher Scientific Cat# Q33231). The sequencing of the libraries was performed at the BGI Oceanic headquarters using MGI DNBSEQ T7 sequencer, with 1260 million reads targeted recovery per sample.

### Bioinformatics Workflow

#### Preprocessing

The stitched .czi files taken with the Z2 Axio Imager were processed through ImageStudio v 2.1.1 using the ssDNA setting for staining to ensure they passed QC. They were then transformed into .ipr files to ensure compatibility with the SAW pipeline. The paired FASTQ, mask and transformed .ipr files were then processed using the SAW pipeline v 6.12 using default parameters including cell binning. Raw FASTQ files generated from mouse hearts were aligned using default SAW mapping parameters to the reference genome GRCm39 while raw FASTQ files generated from organoids were aligned to the reference genome GRCh38.

#### Analysis of organoids and reference tissues

All organoids and in vivo reference tissues was conducted using the Squidpy package v1.2.2 [32] and Stereopy package v1.2 [33] with Python v3.8.2.

#### Evaluation of quality metrics

Quality metrics were evaluated globally across the whole chip for each organoid type, and at the individual-organoid level for organoids of interest. Number of counts and number of genes per chip were computed using Stereopy, while number of mapped reads, mRNA-captured DNA nanoballs under tissue and sequencing saturation values were extracted from SAW output files and report. Per-organoid metrics were computed after filtering the data to retain only a single organoid, using the coordinates extracted with the with the plt.interact_spatial_scatter() function of Stereopy. This analysis was repeated for 3 different bin sizes (50, 75 and 100). Before computing the number of bins, genes and counts, a filtering was applied to remove signal outside of the organoid boundaries and bins with extremely low counts or high mitochondrial gene percentage. This filtering ensured that we counted and evaluated only bins representing true biological signal. The filtering criteria varied between kinds of organoids and tissues as it was highly dependent on the specific counts’ distribution. In some case, poor-quality bins were not limited to clear outliers but represented a substantial part of the distribution, making outlier-based filtering inadequate. The specific filtering steps are documented in the code associated with the publication.

#### Organoid-specific regional analysis to uncover spatially-restricted transcriptomic differences

Artifacts and outliers at the edge of the organoids were first filtered to retain biologically relevant spatial information, as these frequently contained abnormal amount of UMIs counts (Fig. 3A-3C, Fig. S4A-S4C). For each of the brain, cartilage and transwell kidney organoids (which were all largely circular in shape), we first calculated the centroid coordinates, then Euclidean distance of each point from the centroid was computed. We identified points with distances greater than the 90th percentile of all distances as “distant points.” Then we proceeded to detect outliers, that are points located outside the organoid represented false signals. These artifacts needed to be systematically removed to ensure that only biologically relevant spatial information was retained. These artifacts were located at a distance from the rest of the organoid cells, making it necessary to apply a targeted filtering approach. To remove outliers, we applied a k-nearest neighbours (k-NN) approach using the NearestNeighbors() function from sklearn v1.5.0 and ball_tree algorithm parameter. A k parameter of 10 was used for brain organoids and 250 for kidney and cartilage. Since most of the false signals were spatially distant from the organoid cells, k-NN was used to identify points with abnormally high distances to their nearest neighbours. However, to prevent central hollow regions from being misclassified as outliers, we restricted this analysis to the distant bins. For each distant bin, the average distance to its nearest neighbours was computed and standardized using a Z-score transformation. Bins with a Z-score greater than 2 (or higher depending on the specific case) were classified as outliers and subsequently removed from further analysis. (Fig. 3C, Fig. S4D-S4F).

Following the removal of outliers, we computed the convex hull of the remaining bins to define the spatial boundary of the dataset. To further classify points into distinct spatial regions of the ‘edge’, ‘border’ and ‘core’, we calculated the minimum distance of each point to the convex hull boundary. Based on the distance distribution we classified each bin as either belonging to the: ‘edge’ for bins within the 10th percentile of boundary distances, ‘border’ for bins with distances between the edge threshold and half of the maximum distance, and ‘core’ for bins with distances exceeding the border threshold (Fig. 3D, Fig. S4G-S4I). The ‘edge’ of the organoid often contained an abnormal number of counts and was removed from further analysis. After spatial classification, we generated pseudo-bulk samples using core and border regions of individual organoids placed on the same chip and performed differential expression analysis between the core and border regions of the same organoid types. Pseudobulk was performed using the pydeseq2 v1.6.0 and decoupler v0.4.11 packages. The get_pseudobulk() function was used to generate pseudobulk samples from the anndata object using the core and border regions from each kind of organoid. This was then built into a DeSeq2 object and differential expression analysis performed. We treated the organoids on the same chip as pseudo-replicates, as even if the organoids were not exactly biological replicates but belonged to either different lines subjected to editing (brain organoids) or different growing conditions (cartilage organoids), the aim of the analysis was to identify regional differences in general within the inner and outer areas of the organoids, regardless of the organoid conditions. Subsequently, we conducted Gene Set Enrichment Analysis (GSEA) to determine enriched pathways and biological processes associated with differentially expressed genes.

For the heart organoids which had an elliptic shape, to approximate the organoid boundary, a convex hull was computed using the set of spatial coordinates. Specifically, the algorithm Quickhull was used to identify the smallest convex polygon capable of enclosing all points. Then each bin’s distance to the convex hull boundary was computed by treating each bin’s coordinates as a point and measuring its distance to the perimeter of the convex polygon. This metric provided an estimate of whether a bin was located near the tissue edge (small distance) or more centrally (larger distance). To remove perimeter effects and focus on interior tissue regions, bins in the lower 15th percentile of boundary distance were excluded. The remaining data (with distances above this threshold) formed the “filtered” subset of bins, considered representative of the tissue’s interior.

The spatial coordinates of the filtered bins were then subjected to PCA. The first principal component (PC1) was assumed to capture the tissue’s major “long axis.” This vector (PC1) was used for subsequent segmentation of the tissue into poles and centre regions. Each filtered bin was projected onto the long-axis vector by calculating a dot product between the spot’s spatial coordinates and the principal-axis vector. This yielded a one-dimensional scalar value for each spot, referred to as its “projected distance.” The minimum and maximum projected distances defined the overall range (or “length”) along this axis. The axis was then divided into three segments of equal length, the first segment from the minimum projection to one-third of the total range, the middle segment covering the next one-third of the range and the final segment from two-thirds of the range to the maximum projection. Bins in the bottom or top third of this range were labelled as “pole”, whereas spots in the central third were labelled as “centre.” Differential expression analysis in pseudo-bulk was performed between centre of the serum free and directed maturation heart organoids.

## Figure Legends

Supplementary Figure 1 - Overview of data capture across different organoids and in vivo reference tissues.

(A) Total DNA nanoballs captured across each Stereo-seq chip for different organoids and in vivo reference tissues.

(B) Total number of UMIs captured across each Stereo-seq chip for different organoids and in vivo reference tissues.

(C) Sequence depth and sequence saturation curves for different organoids and reference tissues. Vendor recommended 80% saturation cut-off displayed as blue dotted line.

(D) Correlation between total counts and number of CellBins across different organoids passing QC and two in vivo reference tissues.

(E) Total number of CellBins across in vivo reference tissues and individual brain, kidney transwell and cartilage organoids.

(F) Distribution of number of counts per CellBin across in vivo reference tissues and single organoids from chips passing QC.

(G) Distribution of number of genes per CellBin across in vivo reference tissues and single organoids from chips passing QC.

(H) Distribution of number of counts per bin size 50 across in vivo reference tissues and single organoids from chips passing QC.

(I) Distribution of number of genes per bin size 50 across in vivo reference tissues and single organoids from chips passing QC.

Supplementary Figure 2 - Overview of data capture across different organoids and in vivo reference tissues.

(A) Bar plot of percentage of CellBins expression housekeeping genes GAPDH/Gapdh and ACTB/Actb across in vivo reference tissues and single organoids from chips passing QC.

(B) Dot plot of relative intensity of expression of key marker genes across different organoid types and in vivo reference tissues.

(C) Heatmap of gene expression housekeeping and mitochondrial genes across brain, cartilage, heart muscle, kidney transwell and lung organoid chips compared in vivo reference tissues E13.5 mouse head and 21PCW mouse heart.

(D) Heatmap of tissue-specific marker gene expression across brain, cartilage, heart muscle, kidney transwell and lung organoid chips compared in vivo reference tissues E13.5 mouse head and 21PCW mouse heart.

Supplementary Figure 3 – Effect of application of normalisation across chip or per organoid on expression of housekeeping and marker genes for brain, cartilage and kidney transwell organoids.

Supplementary Figure 4 - Analysis of spatially-restricted, differentially expressed genes across centre and border regions for individual brain, cartilage and kidney transwell organoids.

(A) Heatmap of UMI counts across brain organoid 2-3 using CellBin. Regions of high UMI counts are displayed in dark purple.

(B) Heatmap of UMI counts across cartilage organoids 1-4 using CellBin. Regions of high UMI counts are displayed in dark purple.

(C) Heatmap of UMI counts across kidney transwell organoid 1-2 using CellBin. Regions of high UMI counts are displayed in dark purple.

(D) Outlier CellBins identified and visualised in magenta across brain organoid 2-3, representing a false positive signal to be removed before region analysis.

(E) Outlier CellBins identified and visualised in magenta across cartilage organoid 1-4, representing a false positive signal to be removed before region analysis.

(F) Outlier CellBins identified and visualised in magenta across kidney organoid replicates 1-2, representing a false positive signal to be removed before region analysis.

(G) Edge (blue), border (green) and centre (red) regions highlighted for analysis in brain organoid replicates 2-3.

(H) Edge (blue), border (green) and centre (red) regions highlighted for analysis in cartilage organoids 1-4.

(I) Edge (blue), border (green) and centre (red) regions highlighted for analysis in kidney organoids 1-2.

(J) Volcano plots of genes upregulated in core displayed in red and those upregulated in border region visualised in blue for cartilage organoid samples 1-4. The analysis did not return statistically significant differentially expressed genes.

(K) Heatmap of UMI counts 3 selected control heart muscle organoids and 3 selected DM organoids.

(L) Visualisation of centre and poles for 3 selected control heart muscle organoids and 3 selected DM organoids.

Supplementary Table 1 – Overview of number of retained bins and corresponding tissue area across bin size 50, 75 and 100 for in vivo reference tissues E13.5 mouse head and 21 PCW mouse heart and Brain organoid 1 and Cartilage organoid .1

## Resource Availability

### Lead Contact

Further information and requests for resources should be directed to and will be fulfilled by the lead contacts, Fernando Rossello and Mirana Ramialison.

### Data and code availability

- All original code is available on Github: https://github.com/Ramialison-Lab/StereoseqOrganoids

## Acknowledgements

This work was supported by the Novo Nordisk Foundation Center for Stem Cell Medicine, reNEW, supported by Novo Nordisk Foundation grant number NNF21CC0073729; a BGI STOmics Grant to NC and MR. E.R.P. is supported by an Investigator Grant from the National Health and Medical Research Council (NHMRC) of Australia (GNT2008376). S.V. is supported by an Investigator Grant from the NHMRC of Australia (GNT2025776) and a Fellowship from the Sarah and Lachlan Murdoch Foundation. H.K.V was supported by a Postdoctoral Fellowship (106645) from the National Heart Foundation of Australia. MR is funded by a Future Leader Fellowship (107328) from the Heart Foundation of Australia, a Human Frontiers Science Program Grant (RGP008/2024). MR and HTN are supported by an NHMRC Ideas Grant (APP1180905). Additional infrastructure funding to the Murdoch Children’s Research Institute was provided by the Australian Government National Health and Medical Research Council Independent Research Institute Infrastructure Support Scheme. The Australian Regenerative Medicine Institute is supported by grants from the State Government of Victoria and the Australian Government.

## Author contributions

Conceptualization: MN, NC, RFL, MR. Methodology: MN, NC, RFL, MR. Software: MN. Formal Analysis: MN. Investigation: MN, NC, RFL, AL, LR, RS, AM, MS, NT, VS, DT, MW, PXE, SKT, RS, FA, HV, HN. Resources: XS, BY, KLL. Data Curation: MN, NC, RFL. Writing Original Draft: MN, NC, RFL, FJR, MR. Writing review and editing: all authors. Supervision: SV, GB, MCF, DDE, FJR, MR, DAE, ERP, RJM. Funding acquisition: NC, MR, AGE, SRL, KTL, JMV, RJM, ESN, EGS, RBW, MHL, DAE, PER, HKV, MCF, DDE, SV.

---

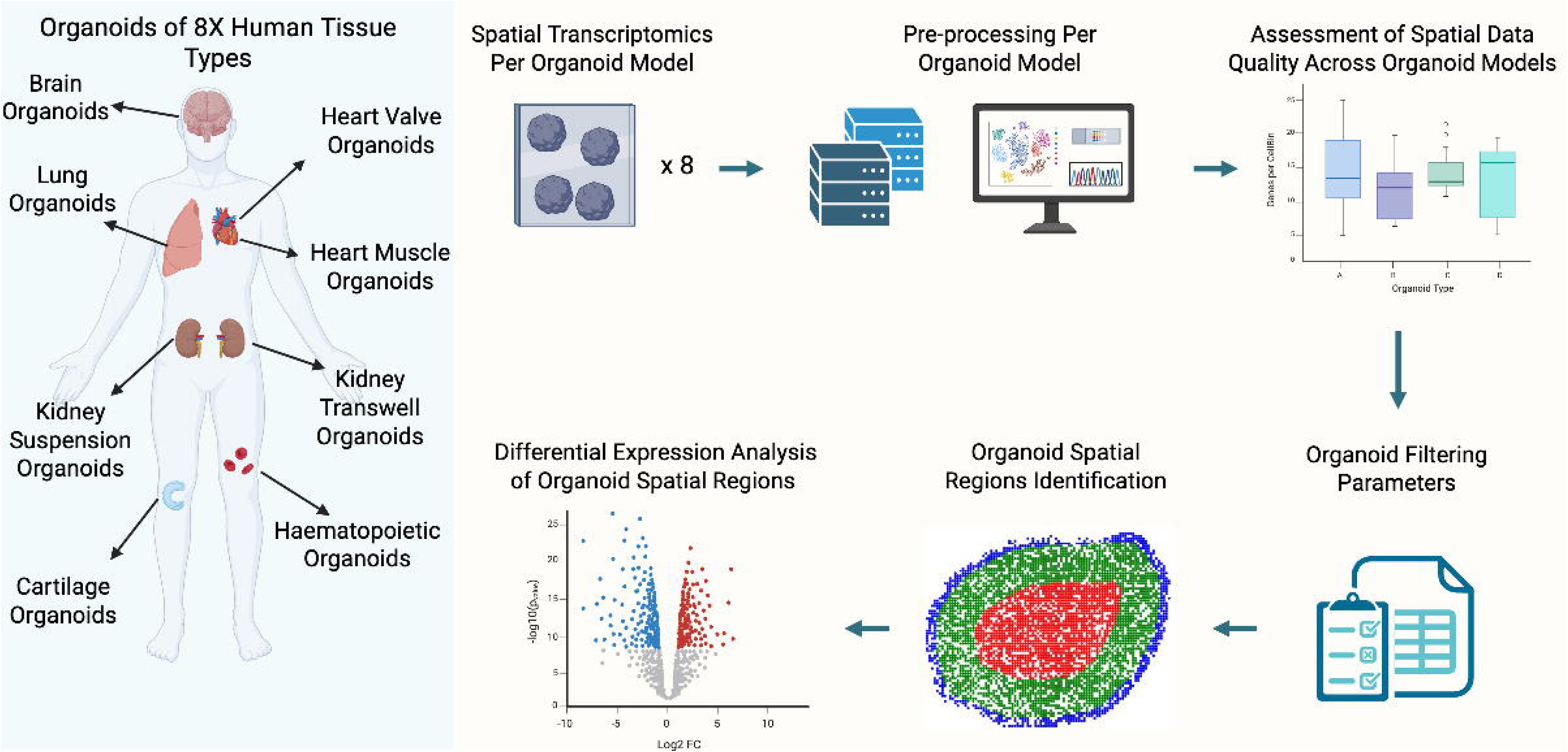

## Bibliography

[1] D. Zhao, W. Lei, and S. Hu, “Cardiac organoid — a promising perspective of preclinical model,” Dec. 01, 2021, BioMed Central Ltd. doi: 10.1186/s13287-021-02340-7.

[2] M. Prasad, R. Kumar, L. Buragohain, A. Kumari, and M. Ghosh, “Organoid Technology: A Reliable Developmental Biology Tool for Organ-Specific Nanotoxicity Evaluation,” Sep. 23, 2021, Frontiers Media S.A. doi: 10.3389/fcell.2021.696668.

[3] Z. Zhao et al., “Organoids,” Nature Reviews Methods Primers, vol. 2, no. 1, Dec. 2022, doi: 10.1038/s43586-022-00174-y.

[4] L. N. Waylen, H. T. Nim, L. G. Martelotto, and M. Ramialison, “From whole-mount to single-cell spatial assessment of gene expression in 3D,” Dec. 01, 2020, Nature Research. doi: 10.1038/s42003-020-01341-1.

[5] A. Chen et al., “Spatiotemporal transcriptomic atlas of mouse organogenesis using DNA nanoball-patterned arrays,” Cell, vol. 185, no. 10, pp. 1777–1792.e21, May 2022, doi: 10.1016/j.cell.2022.04.003.

[6] P. Wahle et al., “Multimodal spatiotemporal phenotyping of human retinal organoid development,” Nat Biotechnol, vol. 41, no. 12, pp. 1765–1775, Dec. 2023, doi: 10.1038/s41587-023-01747-2.

[7] Chiaradia et al., “Tissue morphology influences the temporal program of human brain organoid development,” Cell Stem Cell, vol. 30, no. 10, pp. 1351–1367.e10, Oct. 2023, doi: 10.1016/j.stem.2023.09.003.

[8] S. Ma et al., “Lamination-based organoid spatially resolved transcriptomics technique for primary lung and liver organoid characterization,” Proc Natl Acad Sci U S A, vol. 121, no. 46, Nov. 2024, doi: 10.1073/pnas.2408939121.

[9] Y. Lin et al., “Evaluating stably expressed genes in single cells,” Gigascience, vol. 8, no. 9, Sep. 2019, doi: 10.1093/gigascience/giz106.

[10] I. Hautefort, M. Poletti, D. Papp, and T. Korcsmaros, “Everything You Always Wanted to Know About Organoid-Based Models (and Never Dared to Ask),” Jan. 01, 2022, Elsevier Inc. doi: 10.1016/j.jcmgh.2022.04.012.

[11] S. P. Moss, E. Bakirci, and A. W. Feinberg, “Engineering the 3D structure of organoids,” Jan. 14, 2025, Cell Press. doi: 10.1016/j.stemcr.2024.11.009.

[12] S. R. Lamandé et al., “Modeling human skeletal development using human pluripotent stem cells,” Proc Natl Acad Sci U S A, vol. 120, no. 19, May 2023, doi: 10.1073/pnas.2211510120.

[13] A. Uzquiano et al., “Proper acquisition of cell class identity in organoids allows definition of fate specification programs of the human cerebral cortex,” Cell, vol. 185, no. 20, pp. 3770–3788.e27, Sep. 2022, doi: 10.1016/j.cell.2022.09.010.

[14] R. J. Mills et al., “Functional screening in human cardiac organoids reveals a metabolic mechanism for cardiomyocyte cell cycle arrest,” Proc Natl Acad Sci U S A, vol. 114, no. 40, pp. E8372–E8381, Oct. 2017, doi: 10.1073/pnas.1707316114.

[15] H. K. Voges et al., “Vascular cells improve functionality of human cardiac organoids,” Cell Rep, vol. 42, no. 5, May 2023, doi: 10.1016/j.celrep.2023.112322.

[16] M. Pocock et al., “Maturation of human cardiac organoids enables complex disease modelling and drug discovery,” Sep. 09, 2024. doi: 10.1101/2024.09.05.611336.

[17] P. Xie et al., “Digital reconstruction of full embryos during early mouse organogenesis,” Cell, vol. 188, no. 17, pp. 4754–4772.e18, Aug. 2025, doi: 10.1016/j.cell.2025.05.035.

[18] M. F. Oliveira et al., “Characterization of immune cell populations in the tumor microenvironment of colorectal cancer using high definition spatial profiling,” Jun. 05, 2024. doi: 10.1101/2024.06.04.597233.

[19] Vlahos et al., “Generation of iPSC lines from peripheral blood mononuclear cells from 5 healthy adults,” Stem Cell Res, vol. 34, Jan. 2019, doi: 10.1016/j.scr.2018.101380.

[20] M. Vanslambrouck et al., “A toolbox to characterize human induced pluripotent stem cell-derived kidney cell types and organoids,” Journal of the American Society of Nephrology, vol. 30, no. 10, pp. 1811–1823, 2019, doi: 10.1681/ASN.2019030303.

[21] R. B. Werder et al., “CRISPR interference interrogation of COPD GWAS genes reveals the functional significance of desmoplakin in iPSC-derived alveolar epithelial cells,” Sci Adv 2022. doi: 10.1126/sciadv.abo6566

[22] M. Yammine, S. Mirda Abularach, L. Sampurno, J. F. Bateman, S. R. Lamandé, and M. D. Shoulders, “Using CRISPR/Cas9 to generate a heterozygous COL2A1 p.R719C iPSC line (MCRIi019-A-6) model of human precocious osteoarthritis,” Stem Cell Res, vol. 67, Mar. 2023, doi: 10.1016/j.scr.2023.103020.

[23] M. Abo et al., “Air-liquid interface culture promotes maturation and allows environmental exposure of pluripotent stem cell-derived alveolar epithelium,” Resource and Technical Advance 2022, doi: 10.1172/jci.insight.155589.

[24] F. J. Hawkins et al., “Derivation of airway basal stem cells from human pluripotent stem cells,” Cell Stem Cell, vol. 28, no. 1, pp. 79–95.e8, 2021, doi: 10.1016/j.stem.2020.09.017.

[25] A. Jacob et al., “Differentiation of Human Pluripotent Stem Cells into Functional Lung Alveolar Epithelial Cells,” Cell Stem Cell, vol. 21, no. 4, pp. 472–488.e10, Oct. 2017, doi: 10.1016/j.stem.2017.08.014.

[26] E. S. Ng et al., “Long-term engrafting multilineage hematopoietic cells differentiated from human induced pluripotent stem cells,” Nat Biotechnol, 2024, doi: 10.1038/s41587-024-02360-7.

[27] H. K. Voges, R. J. Mills, E. R. Porrello, and J. E. Hudson, “Generation of vascularized human cardiac organoids for 3D in vitro modeling,” STAR Protoc, vol. 4, no. 3, Sep. 2023, doi: 10.1016/j.xpro.2023.102371.

[28] T. Lawlor et al., “Cellular extrusion bioprinting improves kidney organoid reproducibility and conformation. HHS Public Access,” Nat Mater, vol. 20, no. 2, pp. 260–271, 2021, doi: 10.6084/m9.figshare.12957122.

[29] S. V. Kumar et al., “Kidney micro-organoids in suspension culture as a scalable source of human pluripotent stem cell-derived kidney cells,” Development (Cambridge), vol. 146, no. 5, Mar. 2019, doi: 10.1242/dev.172361.

[30] S. Velasco et al., “Individual brain organoids reproducibly form cell diversity of the human cerebral cortex,” Nature, vol. 570, no. 7762, pp. 523–527, Jun. 2019, doi: 10.1038/s41586-019-1289-x.

[31] J. Huang et al., “SARS-CoV-2 Infection of Pluripotent Stem Cell-Derived Human Lung Alveolar Type 2 Cells Elicits a Rapid Epithelial-Intrinsic Inflammatory Response,” Cell Stem Cell, vol. 27, no. 6, pp. 962–973.e7, Dec. 2020, doi: 10.1016/j.stem.2020.09.013.

[32] G. Palla et al., “Squidpy: a scalable framework for spatial omics analysis,” Nat Methods, vol. 19, no. 2, pp. 171–178, Feb. 2022, doi: 10.1038/s41592-021-01358-2.

[33] S. Fang et al., “Stereopy: modeling comparative and spatiotemporal cellular heterogeneity via multi-sample spatial transcriptomics,” Dec. 05, 2023. doi: 10.1101/2023.12.04.569485.

