## SupplementaryFigures for "Application of spatial transcriptomics across organoids: a high-resolution spatial whole-transcriptome benchmarking dataset"

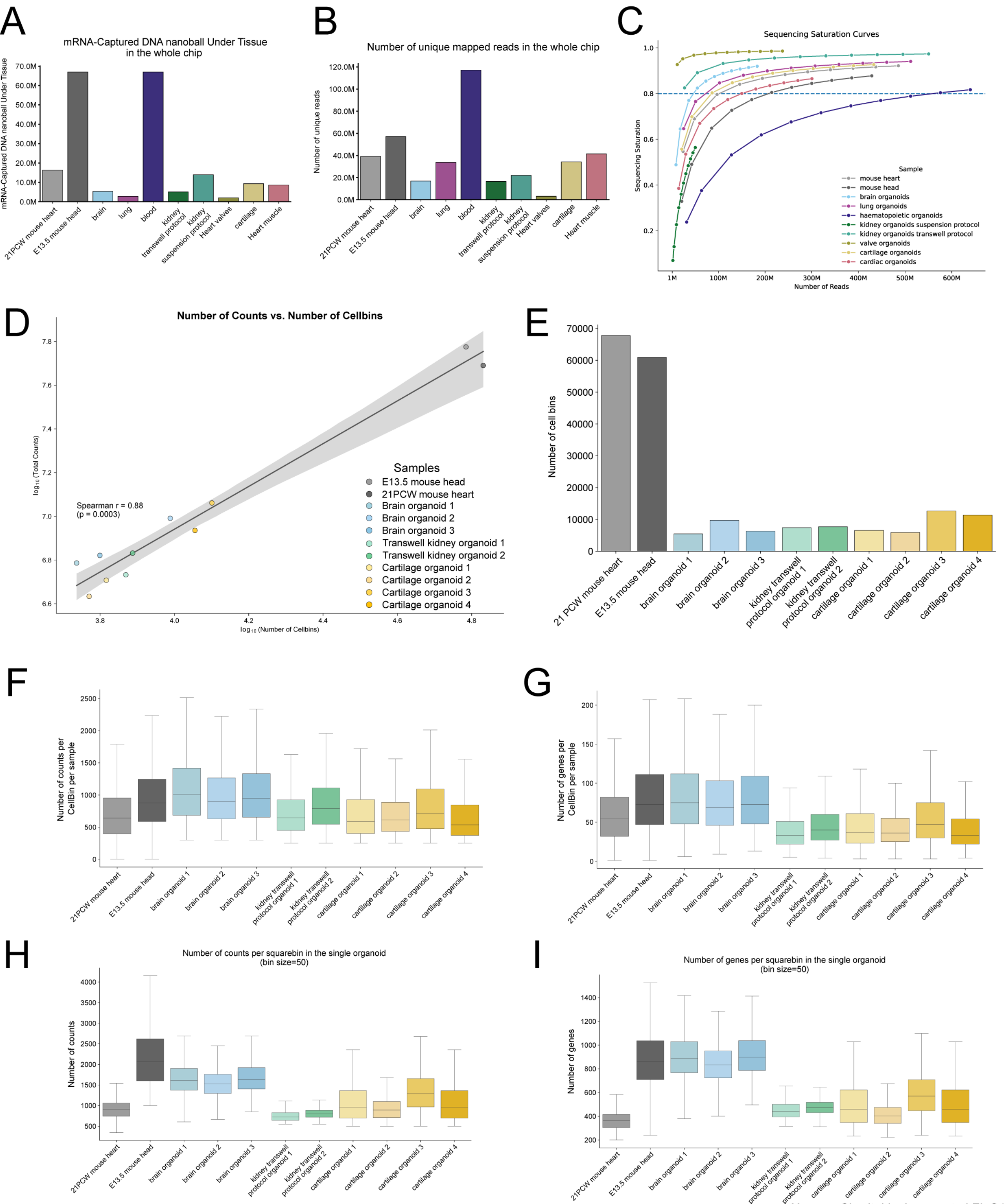


Nucera, Charitakis, Leung et al FigS1

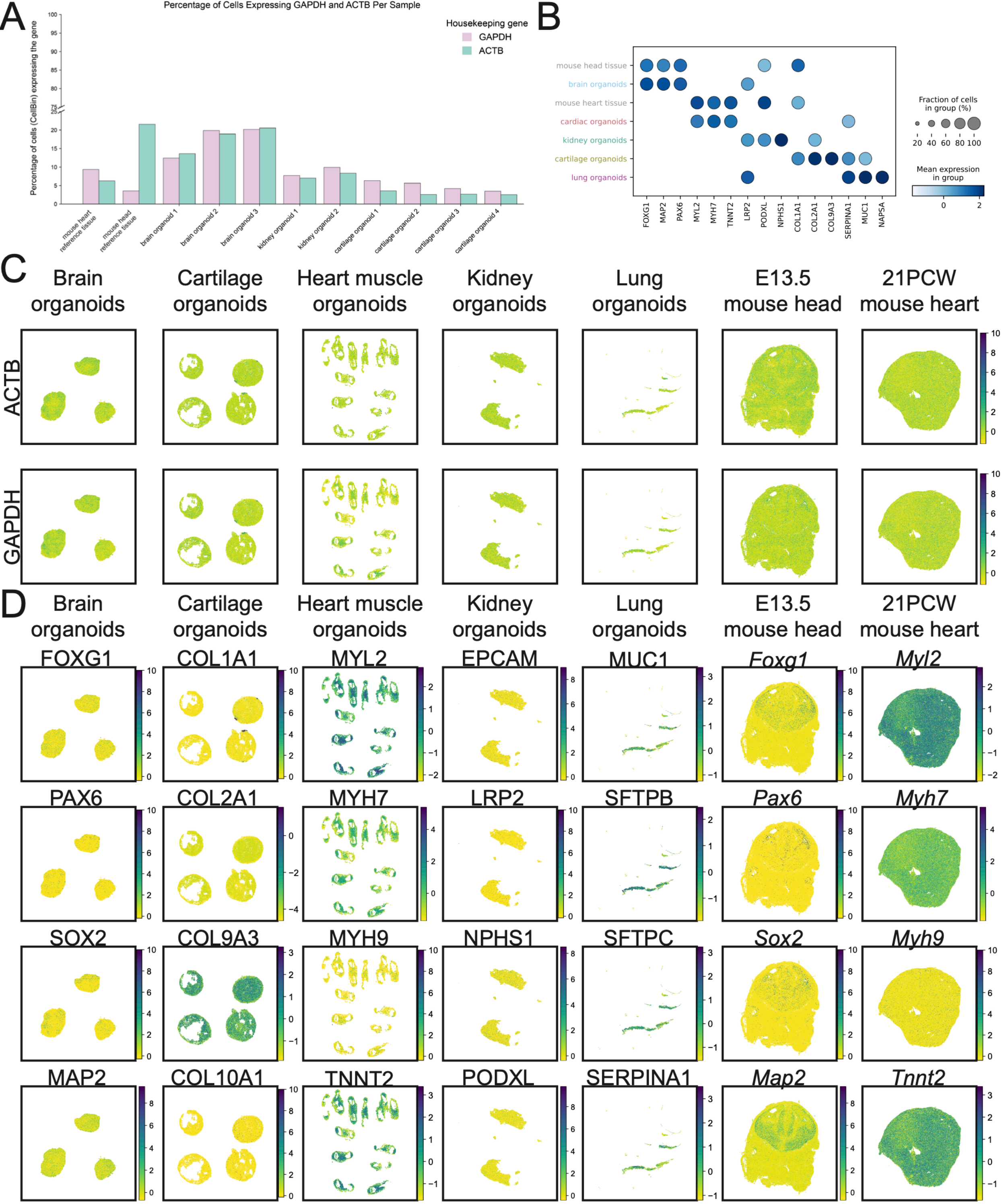


Nucera, Charitakis, Leung et al Fig S2

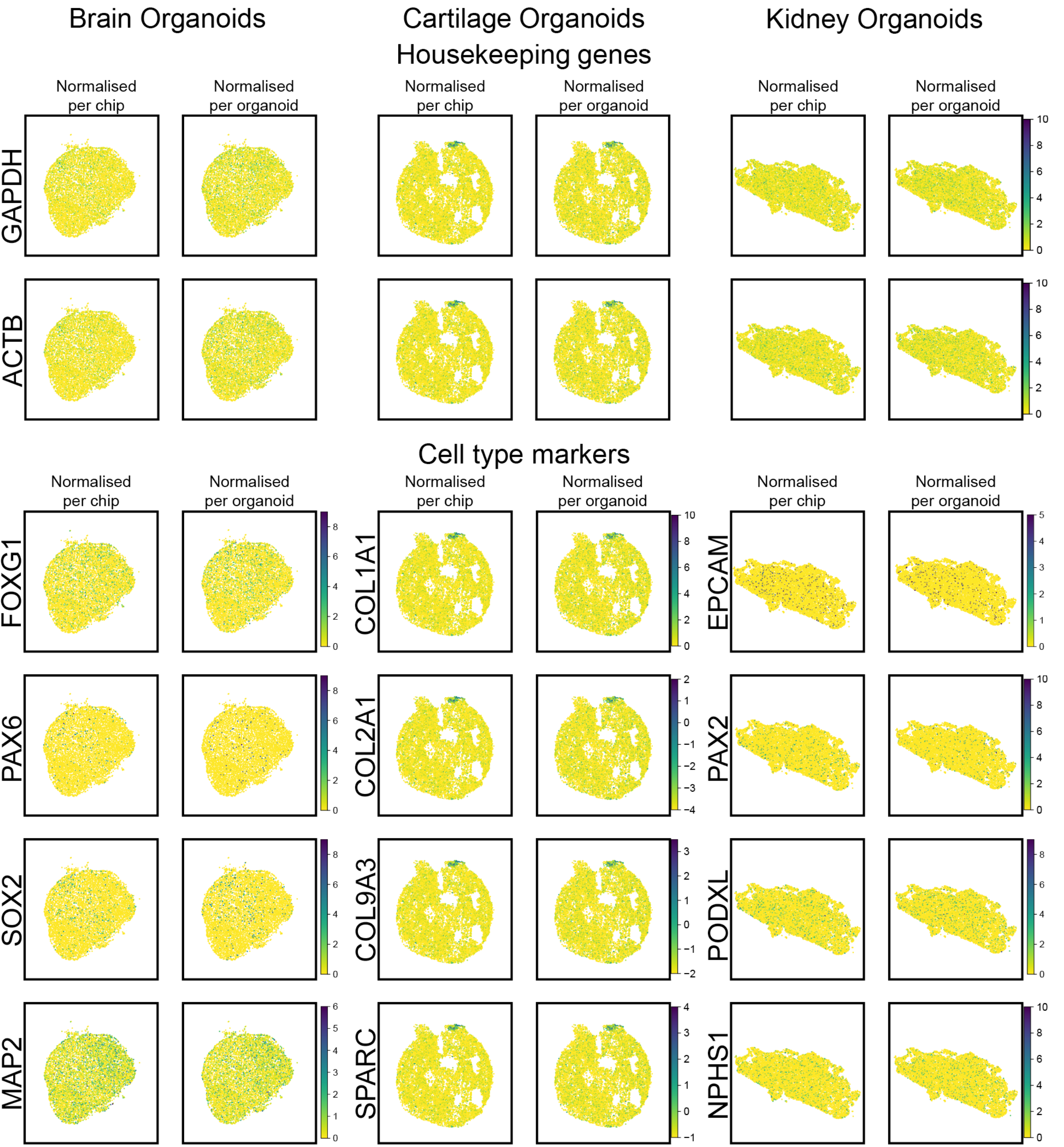


Nucera, Charitakis, Leung et al FigS3

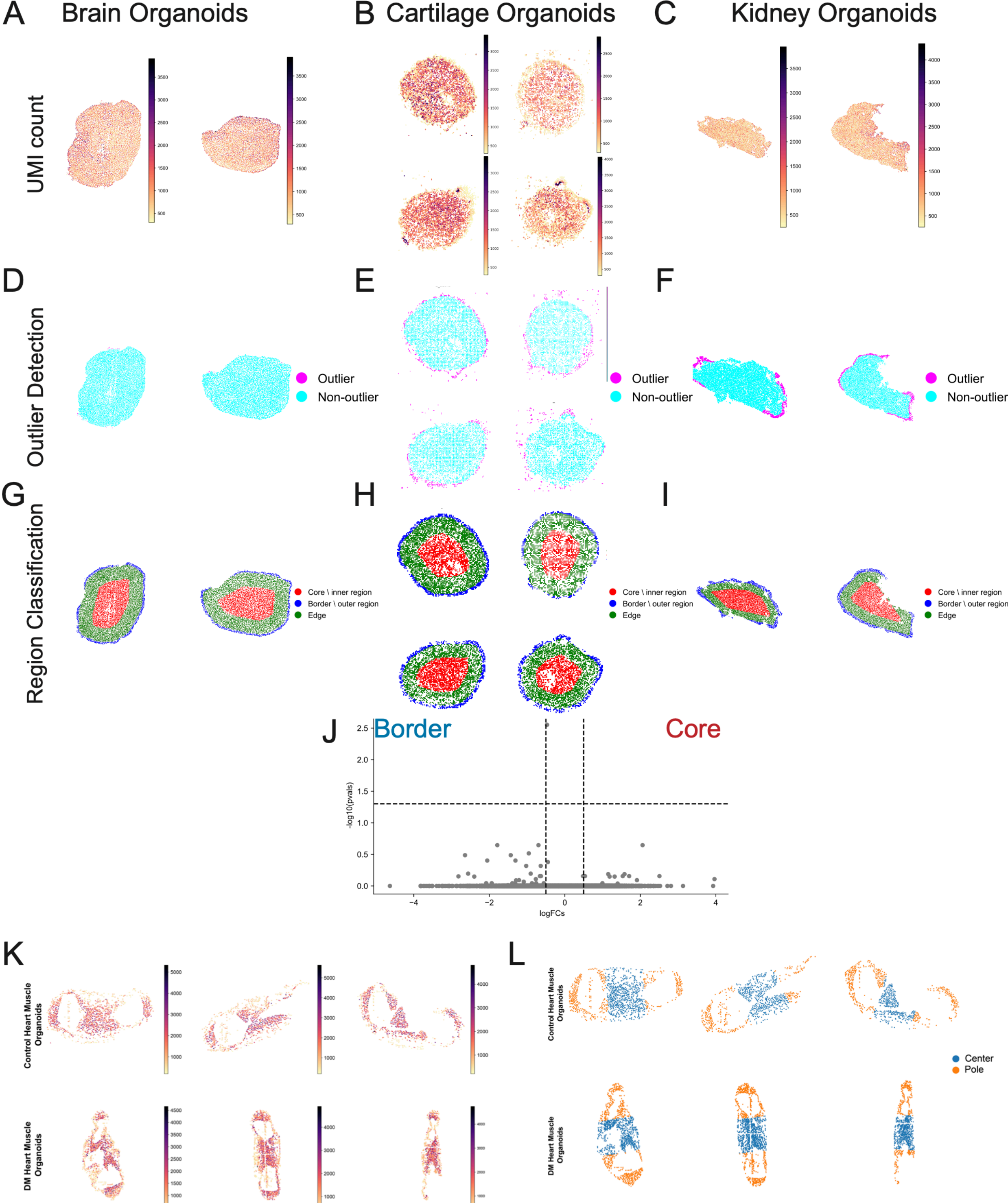


Nucera, Charitakis, Leung et al Fig S4
